# Strain-level and phenotypic stability contrasts with plasmid and phage variability in water kefir communities

**DOI:** 10.1101/2025.02.27.640646

**Authors:** Vincent Somerville, Aiswarya Prasad, Corinne F Maurice, Daniel Garrido-Sanz, Eric Ulrich, Florent Mazel, Garance Sarton-Lohéac, Germán Bonilla-Rosso, Gilles Baud, Jeanne Tamarelle, Lucie Kesner, Malick Ndiaye, Sylvain Moineau, Vincent de Bakker, Vladimir Sentchilo, Yassine El Chazli, SAGE course students 21/22 and 22/23, Philipp Engel

**Affiliations:** University of Lausanne, Switzerland; Agroscope, Liebefeld, Switzerland; Université Laval, Canada; McGill, Canada

**Author notes:** Complete list in acknowledgements (SAGE course students 21/22 and 22/23).

## Abstract

Microbial communities can change in response to top-down factors, such as phages, and bottom-up factors, such as nutrient availability. Previous studies have successfully investigated bacterial species-level dynamics, but diversity and interactions beyond the species-level is usually lacking. Traditional fermented foods, such as water kefir, provide ideal systems to study ecological and evolutionary dynamics beyond the species-level, as they are simple and trackable systems that are cultivated in non-sterile, nutrient-rich environments which foster microbial growth and invasion. Despite the central role of only a few lactic acid bacteria for fermentation, little is known about the genomic diversity and dynamics of these community members over time. Within the framework of a graduate course, 35 students propagated water kefir across several generations under different nutrient conditions and in different households to study microbial responses over time. We found that water kefir communities were generally stable at the species-level, with only rare bacterial species replaced over long timescales (more than 2 years). While we observed little strain-level diversity with few strain replacements over long timescales, closely related strains exhibited variation in accessory gene content, often encoded on plasmids, particularly those involved in ecologically meaningful functions such as sugar utilization pathways and phage defense systems. We hypothesise that these genomic variations could reflect the adaptations of strains to different sugars and phages. Consistent with this, we observed a diverse array of phages, many likely originating from the unique household environments. By documenting the genomic landscape of microbial species, strains, plasmids, and phages, this study advances our understanding of the diversity and dynamics of microbial communities in fermented foods. Furthermore, our course material is publicly available and offers a blueprint for bridging the gap between teaching and research, inspiring the next generation of scientists to unravel the complexities of microbial ecosystems.

**Graphical abstract:** 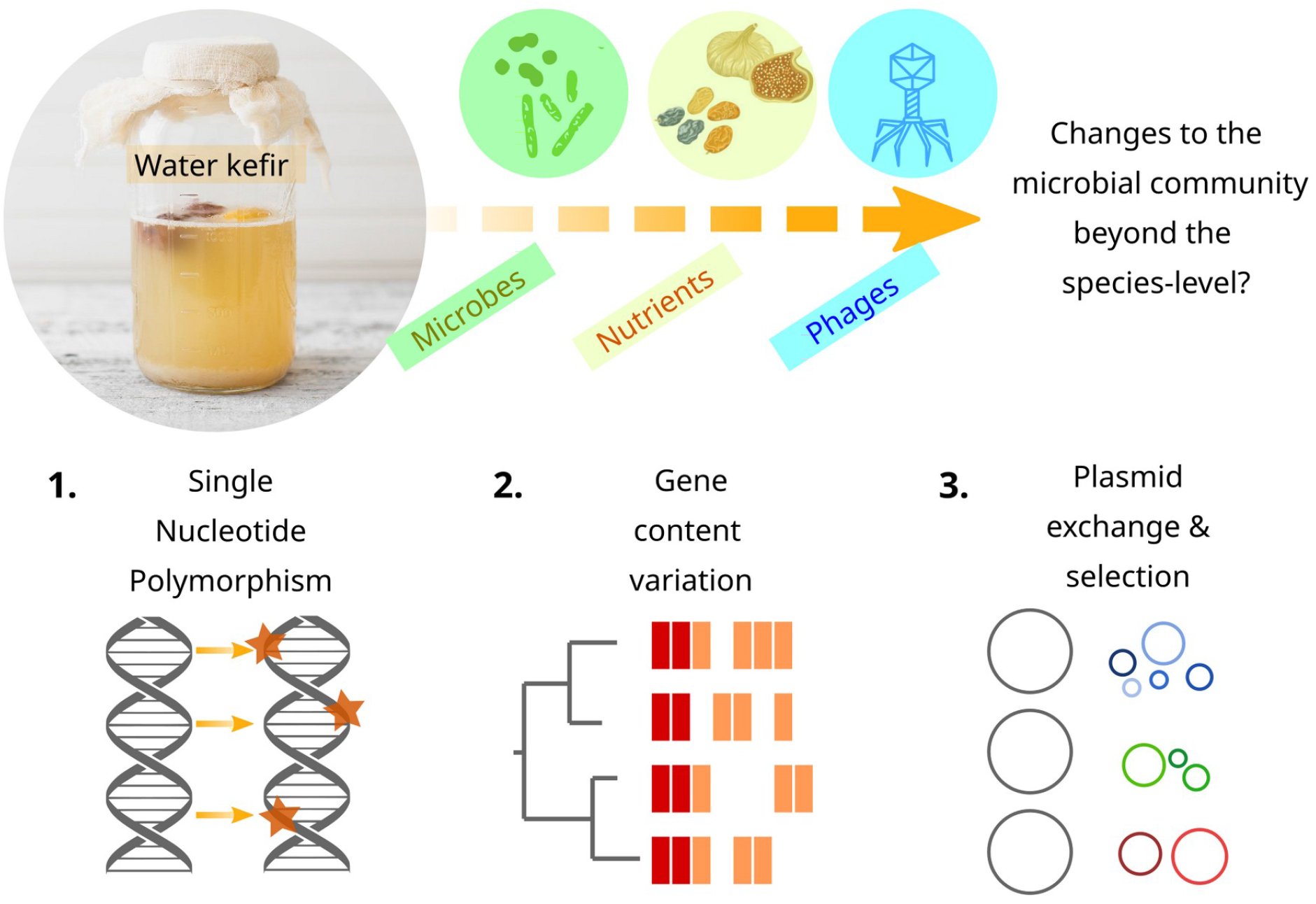

## Introduction

Major ecosystem functions, such as biogeochemical cycles (Arrigo 2005) or the homeostasis of the human gut (Turnbaugh et al. 2007), depend on microbial communities. While these functions are often reliable and reproducible over time, the composition of the underlying microbial communities can vary substantially, even at a relatively coarse taxonomic level like the species- or genus-level (Louca et al. 2016), due to ecological factors such as nutrient availability (Ma et al. 2019), dispersal limitation (Fu et al. 2020), or inter-species interactions (Micali et al. 2023). However, the ecological and evolutionary dynamics within species (*i*.*e*., at the strain-level) remain poorly understood (Van Rossum et al. 2020; Zlitni et al. 2020). This is surprising because functions often differ between strains within species, e.g. in terms of phage resistance (Hussain 2020; Somerville, Schowing, et al. 2022), pathogenicity (Leimbach, Hacker, and Dobrindt 2013) or carbohydrate utilisation (Strachan et al. 2023), with large impacts on ecosystem processes and functions (Kung, Ozer, and Hauser 2010; Shapiro and Polz 2014). There is a need for more studies documenting strain-level diversity patterns across microbial communities and studying the underlying ecological (e.g. niche differentiation, dispersal (Mukherjee et al. 2023; Garud et al. 2019) and selection, drift, speciation and dispersal (Vellend 2010)) and evolutionary (e.g. mutation, drift, and selection (Lenski 2017)) processes.

Traditional fermented foods represent interesting systems to study microbial diversity beyond the species-level for at least three reasons. First, they are stable in composition and reliable in terms of their phenotypic properties (Gänzle 2022). Second, they are usually less diverse than those found in the wild making them more trackable at the strain-level (Louw et al. 2023). Third, they are amenable to experimental manipulation of the environmental settings (e.g. carbon and nitrogen sources). Water kefir is a fizzy beverage created by fermenting sugar water (carbon source) with fruits (nitrogen source) through the transfer and reuse of macroscopic grains called water kefir or tibicos, which contain microorganisms aggregated in dextran exopolysaccharide (Lynch et al. 2021; Martínez-Torres et al. 2017). The water kefir grains contain a microbiota composed of yeast and bacteria (Michielsen et al. 2024). The yeast species, mainly *Saccharomyces cerevisiae* and *Dekkera bruxellensis*, are prevalent members of the community that are important for flavour development (Eckel & Vogel, 2020; Laureys, Cnockaert, De Vuyst, & Vandamme, 2016). The bacterial species encompass several lactic acid bacteria, including *Liquorilactobacillus* spp., *Lentilactobacillus parabuchneri/hilgardii* and *Lacticaseibacillus paracasei* (Fels et al., 2018), as well various acetic acid bacteria and bifidobacteria (Laureys et al. 2018). Although the species composition of water kefir communities varies significantly across kefirs from different origins (Hsieh et al.,2012; Marsh et al., 2013; da Miguel et al., 2011), the fermentation process itself remains phenotypically robust and can be repeated indefinitely by reusing the water kefir grains (Azi et al. 2020; Breselge et al. 2024). Despite this robustness, little is known about how these microbial communities diverge over time, particularly in response to the diverse household environments in which they are cultivated and/or the nutrients provided. To fully understand these dynamics, it is crucial to investigate not only species dynamics but also changes in diversity beyond the species-level. Strain-level diversity in water kefir has primarily been studied through the genomic analysis of isolated strains (Laureys et al. 2016) or metagenome assembled genomes (Verce, De Vuyst, and Weckx 2019). However, less is known about the ecological dynamics, such as strain or plasmid turnover, and evolutionary processes, including *de novo* mutations, that occur over time. In particular, how these changes are influenced by top-down factors, such as phages, or bottom-up factors, such as nutrient availability, remains poorly understood.

In this study, we investigate the dynamics in diversity and microbial composition in water kefir communities across short (weeks) and long (years) timescales and in response to two different environmental conditions (different nitrogen sources). This research was conducted as part of the SAGE (Sequence A GEnome) master’s degree course at the University of Lausanne. The students propagated two closely related water kefir cultures, which had been split approximately two years earlier, using two distinct nutrient resources at home for five passages. By combining sequencing, culturing, and phenotypic analyses, we then explored how microbial communities changed over time. Our findings reveal that i) microbial communities are remarkably stable at the species-level, with only minor changes observed over long timescales (~2 years), ii) strain-level diversity within and between samples is low, iii) gene content variation within species primarily stems from a few plasmids, which encode ecologically important functions, and iv) diverse phages were found across samples.

## Results

### Species-level stability maintained over short and larger also long timescale

We aimed to document microbiota changes in water kefir communities over two timescales. Specifically, to understand the microbial dynamics on a long (evolutionary) timescale, we sampled from two water kefir inocula (start KefirV vs. start KefirP) which share a common origin approximately two years before the start of the experiment (Fig. 1A). To understand the microbial dynamics on a short (ecological) timescale, 35 graduate students at the University of Lausanne grew water kefir, either from KefirV or KefirP, in response to one of two nitrogen sources (figs or raisins) for five successive passages of 36 hours each (2 inocula x 2 nitrogen sources = 4 treatments, Fig. 1A). The amount of grains was measured after each passage, and grains were collected for microbial analysis at the end of the five passages. The grain biomass consistently increased on average by 43% per passage (sd=10%, Supplement Fig. 1), and we detected 2.4*10^7^ CFU (sd=1.2*107, Supplement Fig. 1) per gram of kefir grains. There was no difference between treatments, neither in grain biomass increase nor in bacterial biomass (ANOVA, p-values>0.05).

**Figure 1.**
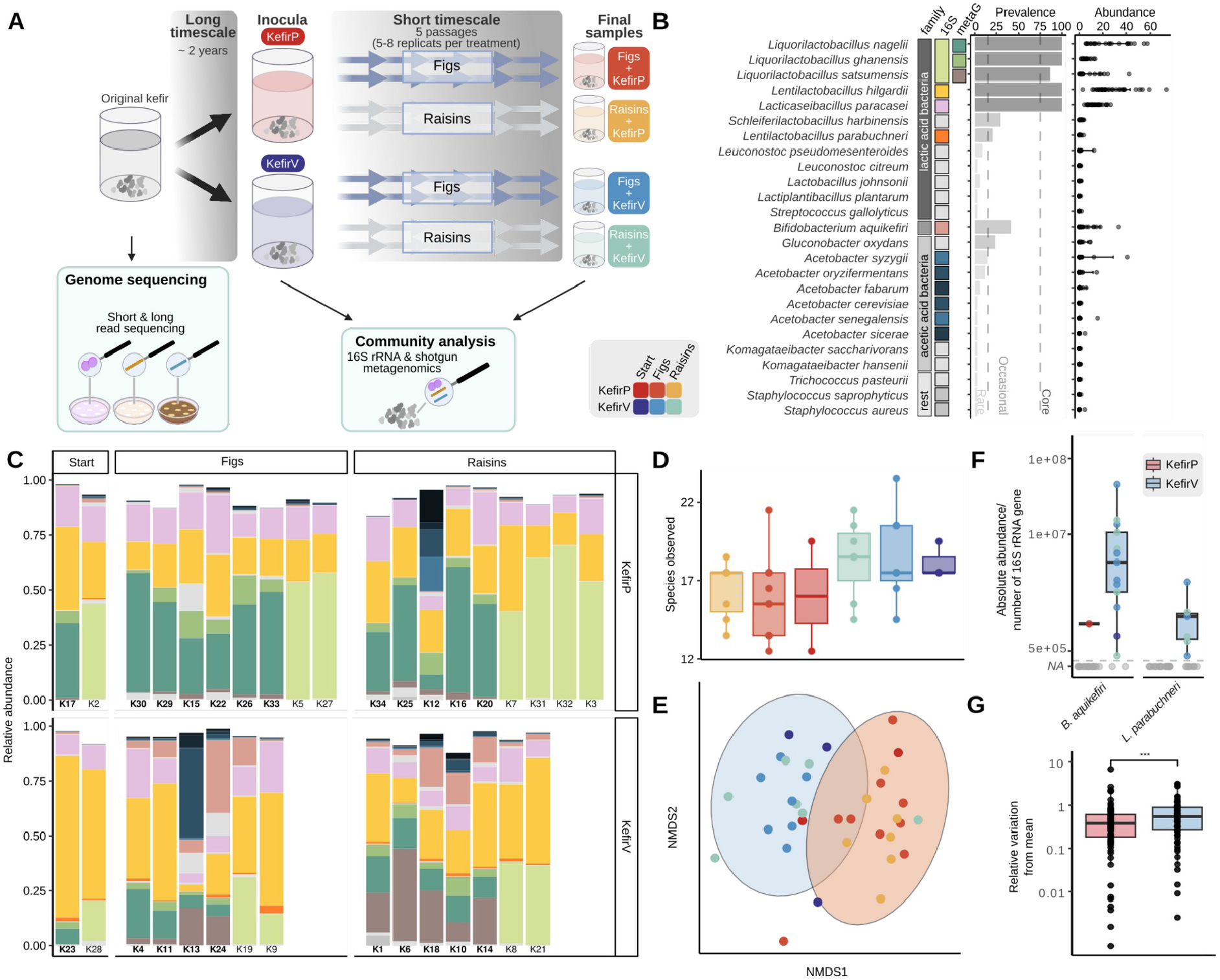
Species diversity in water kefir. A) Experimental design to distinguish between changes on a short and long timescale in water kefir communities. The different treatments and corresponding color scheme, which is used throughout the manuscript, are illustrated in the legend. Additionally, the different sampling time points and methods are illustrated in boxes. B) The prevalence and abundance of the 25 bacterial species, including the categorization of rare, occasional, or core species. C) The abundance of the different bacterial species in the individual samples sorted according to the inoculum vs. nitrogen source (legend corresponds to section B). The samples which were analysed by metagenome sequencing are depicted in bold. The three *Liquorilactobacillus* species are indistinguishable at the 16S rRNA region sequenced, and they were only differentiated with the metagenomic analysis. D) The number of observed species measured in the different water kefir treatments (legend as in Fig. 1A). E) The beta-diversity is illustrated with an NMDS ordination plot of the Bray-Curtis distances where samples are coloured according to treatment. The ellipses illustrate the significant distinguishing factor inoculum (PERMANOVA permutation test, p-value<0.01). F) The absolute abundance of the core and occasional species that only occurred in KefirV. G) The relative deviation of all species in all samples for KefirP vs. KefirV (Wilcoxon test, p-value<0.01).

To analyze the overall composition of the microbial communities, we used partial (V4) 16S rRNA amplicon sequencing (n=35) and shotgun metagenomics on a subset of samples (n=22). To determine the overall microbial composition of our samples, the metagenomic reads were mapped against a defined set of marker genes using mOTU3 (Ruscheweyh et al. 2022). We found that the microbial communities were dominated by bacteria (mean=76%, sd=10%), followed by eukaryotes (mean=23%, sd=10%), and then viruses (~1%; Supplement Fig. 2). In the eukaryotic fraction, we detected two yeast species: *Saccharomyces cerevisiae* (mean=23%, sd=10%) and *Brettanomyces bruxellensis* (mean=1%, sd=1%, Supplement Fig. 2). The 16S rRNA analysis revealed that the bacterial fraction comprised 25 species above a threshold of 0.1% relative abundance, which collectively accounted for 94% of the amplicon sequence variant (ASV) diversity (Fig. 1B/C; Supplement Fig. 3). Five species were prevalent in more than 75% of the samples, and hence we refer to them as the core species for the rest of this study: *Liquorilactobacillus nagelii, Liquorilactobacillus satsumensis* and *Liquorilactobacillus ghanensis* as well as *Lentilactobacillus hilgardii* and *Lacticaseibacillus paracasei* (Fig. 1B/C). Moreover, we detected four species with intermediate prevalence (20-75% of the samples), namely *Bifidobacterium aquikefiri, Gluconobacter oxydans, Lentilactobacillus parabuchneri*, and *Schleiferilactobacillus harbinensis*. Several other species were only present in a few samples (prevalence <20%), some of them belonging to lactic acid bacteria and acetic acid bacteria. For subsequent analyses, we focus exclusively on the bacterial fraction.

To test if bacterial diversity differed across passages and treatments, we carried out alpha- and beta-diversity analysis using the 16S rRNA data. Firstly, the species richness (alpha-diversity) was on average 18 (sd=3) with no significant differences between the samples from the two inocula or the two nutrient conditions (ANOVA, p-value=0.1, Fig. 1D, Supplement Fig. 4). Secondly, microbiota composition, as assessed by beta-diversity analysis using Bray-Curtis distances, clearly clustered by inocula (KefirV vs. KefirP) (PERMANOVA permutation test, p-value<0.01, R^2^=0.2, Fig. 1E). However, the fig and raisin treatments did not have a systematic differential effect on the microbial species composition (PERMANOVA permutation test, p-value=0.3, Fig. 1E).

To pin down which bacterial species differed between the two inocula, we compared absolute species abundance (normalized by 16S qPCR, Wilcoxon-test, p-value<0.01). Three species had similar levels in both inocula (*L. hilgardii, L. satsumensis*, and *G. oxydans*, Supplement Fig. 5), four species were slightly more abundant in the final KefirP sample (*L. nagelli, L. ghanensis, S. harbinensis*, and *L. paracasei*, Supplement Fig. 6), and two species were only present in the final KefirV samples (*B. aquikefiri* and *L. parabuchneri*, Fig. 1F). Two samples (K12, K13) from different treatments stood out as they had very high amounts of acetic acid bacteria (Fig. 1 B/C). Overall, the abundance of the different species was more variable in the final samples for KefirV than for KefirP (Fig. 1G).

### Strain-level stability across short timescales with little strain replacement over long timescale

We showed that species composition was stable across the experiment, but different strains from the same species may be varying between samples, or multiple strains could co-occur in the same sample. To detect such strain-level diversity, we mapped the metagenomic reads (n=22 samples) against a non-redundant genome database consisting of representative bacteria and yeasts from NCBI and isolates identified in our community analysis (see above). We then counted the number of single nucleotide polymorphisms (SNPs) in the mapped reads as a proxy for strain-level diversity within samples. To enable comparison across samples, the analysis was restricted to the five most prevalent species (i.e. the core species), which had a mean coverage over 126x (std = 365x, Supplement Fig. 8). We found between 0.9 and 0.001 SNPs per 100 bps depending on the species and the kefir culture (Fig. 2A). Most of the variation in strain-level diversity was explained by the species, where *L. nagelli* and *L. satsumensis* (> 0.1 SNP per 100 bps) had much higher strain-level diversity than *L. ghanensis* and *L. paracasei* (> 0.01 SNP per 100 bps, ANOVA, p-value<0.05). Almost no strain-level diversity was observed for *L. hilgardii* (0.001 SNP per 100 bps). Moreover, in all samples derived from the KefirP culture we detected between 10-1000x more SNPs than in those from KefirV (Fig. 2A). Strain-level diversity below 1% is generally regarded as low and is typical for natural communities used in food fermentation (Somerville, Berthoud, et al. 2022). This is much lower than for species found in other environments, such as in the gut (Schloissnig et al. 2013), where species generally have > 1% variable sites.

**Figure 2.**
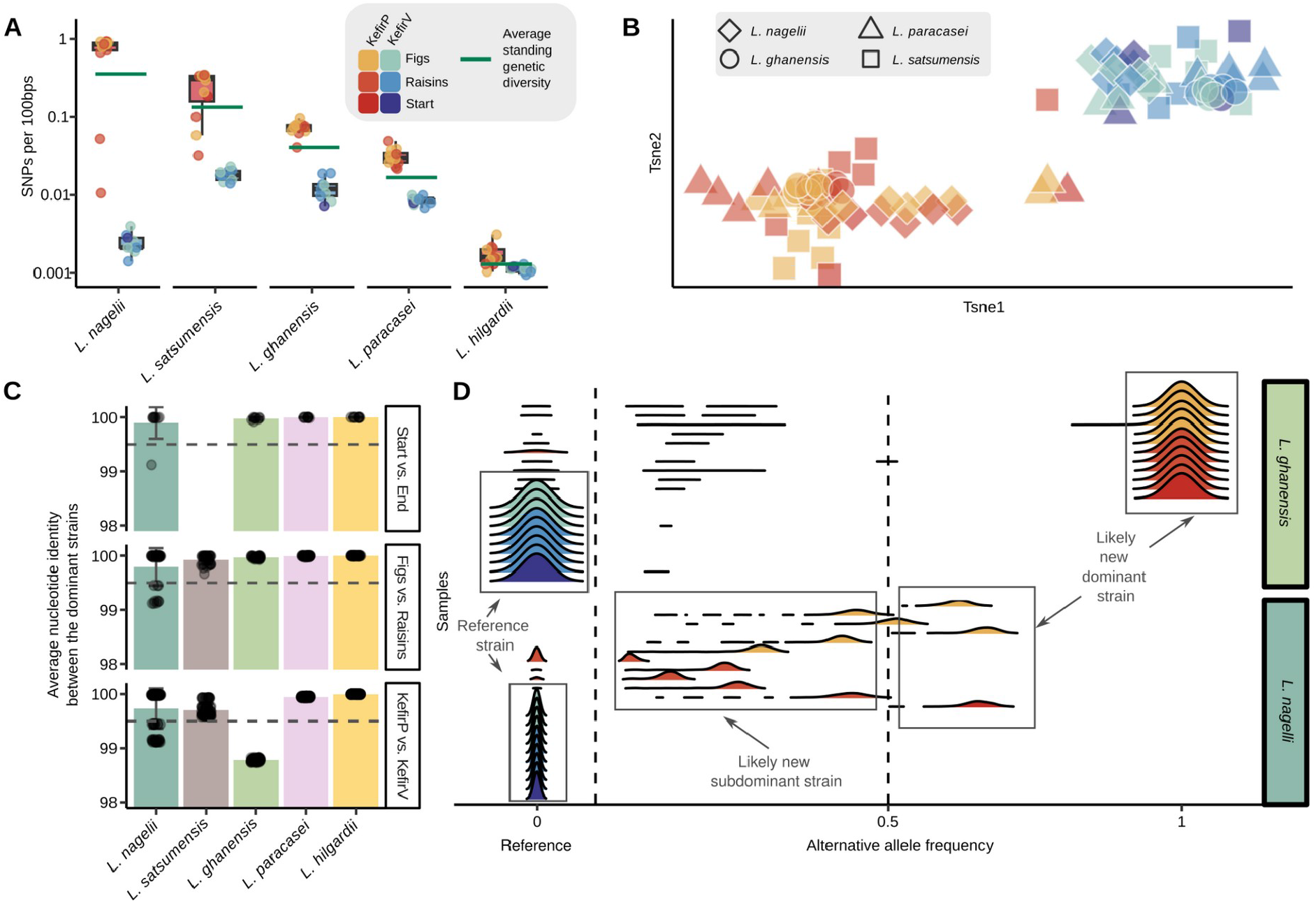
Single nucleotide diversity within and between samples. A) Standing genetic diversity measured by the number of SNPs detected per 100 base pairs (bps) within the different samples split according to inoculum and species. The green line indicates the species average. B) The difference in strain composition between the different samples illustrated in a t-SNE (different species have different shapes, colors correspond to legend in A). Independent t-SNE carried out for each species have been overlaid C) The comparison of the ANI of the dominant alleles for the different start vs. end (top), nitrogen sources (middle) and the two inocula (bottom). D) Difference between the variation in strain frequencies compared to the occurrence of different strains with alternative allele frequencies at all SNP loci of *L. ghanensis* (top) and *L. nagelli* (bottom). Each row corresponds to a sample and a given species, and the distribution of SNP alternative (=non-dominant) allele frequency is plotted.

To test if strain composition differed between the treatment groups, i.e. nutrient condition, or inoculum (KefirP vs KefirV), we carried out beta-diversity analysis. For the three *Liquorilactobacillus* species (*L. nagelii, L. satsumensis, L. ghanensis*) and *L. paracasei*, samples from the same inoculum (PERMANOVA, p<0.01) were more similar to each other in composition, but not from the same nitrogen source (PERMANOVA, p>0.01; Fig. 2B). Notably, the fifth core member, *L. hilgardii*, did not show any clustering, probably due to the low nucleotide diversity present in this species.

To identify if there are different co-existing strains, we estimated average nucleotide identity (estANI, (Olm et al. 2021)) by considering the dominant metagenomic SNPs (allele frequency>0.5) between sample pairs, normalized by the total number of shared loci. For the majority of comparisons the estANI was well above 99.5% (Van Rossum et al. 2020) (Supplement Fig. 9), suggesting that the same (or very similar) strains are present across samples, independent of the nutrient condition, the inoculum, or the sampling time point (Fig. 2C). The only two notable exceptions were *L. ghanensis* and *L. nagelii* for which distinct strains were found between the two different inocula (Fig. 2D).

### Ecologically relevant genes change between closely related strains

Although little strain-level diversity was detected within species in the water kefir cultures, even closely related strains can vary in gene content which may translate into functional differences. To assess differences in gene content between strains of the same species, we isolated, sequenced, and assembled 14 circular genomes from five species. Moreover, we assembled 129 good quality (completeness >85% and contamination <5%) metagenome-assembled genomes (MAGs) from 19 bacterial species (Fig. 3A).

**Figure 3.**
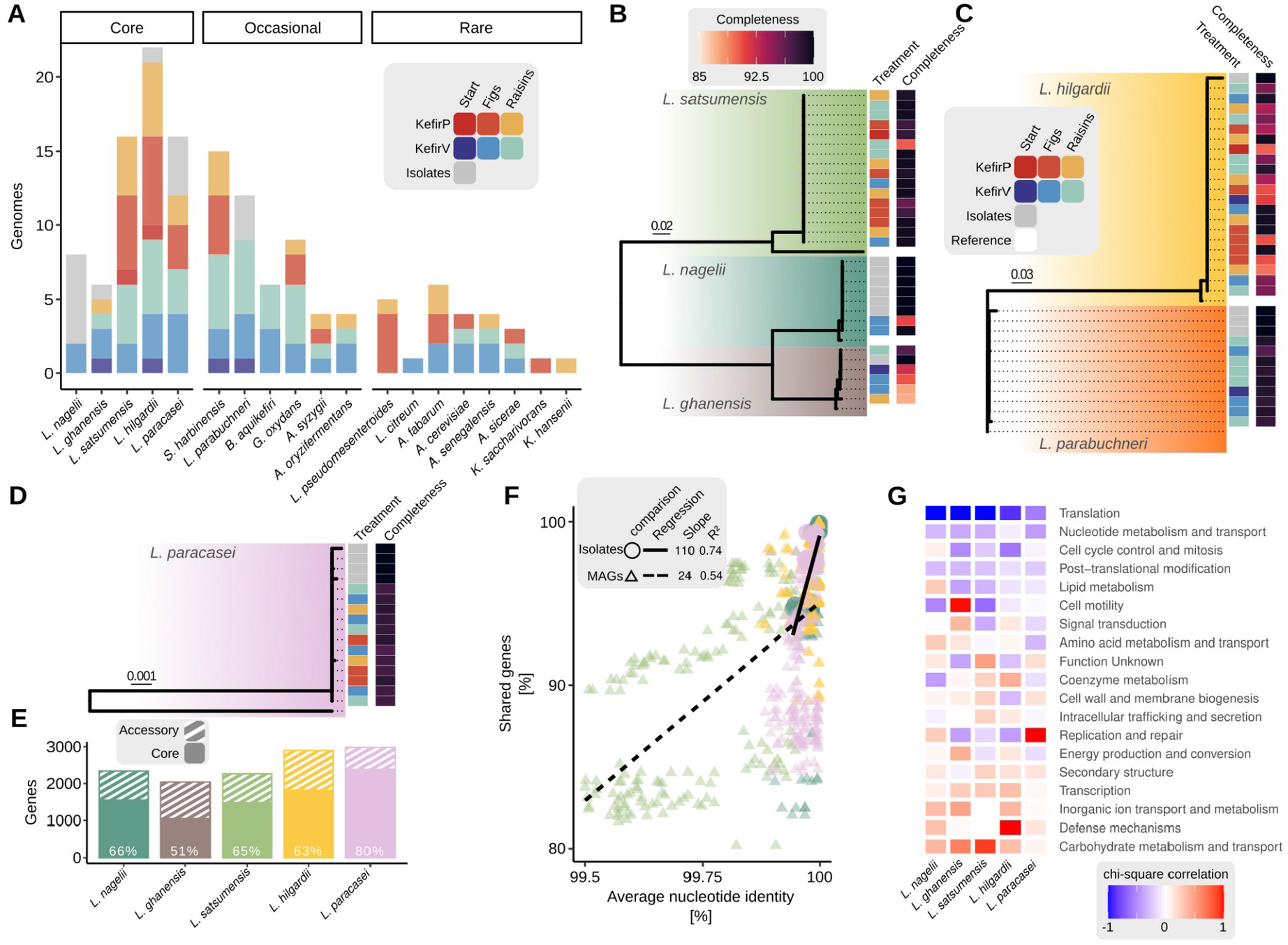
Gene content diversity of the genome collection. A) Sequenced isolates and metagenome-assembled genomes (MAGs) from different species of the core as well as occasional or rare species. B)C)D) The genome phylogenies of the three genera (*Liquorilactobacillus, Lentilactobacillus, and Lacticaseibacillus*) which encompass the core species including all isolate genomes and MAGs and one reference strain per species. The completeness of each MAG and the treatment are noted in the adjacent heatmap. E) The size of the core and accessory genomes in the genome collection. F) The decrease in shared genes with decreasing phylogenetic distance for the MAGs (dashed line) and isolate genomes (line). G) The enrichment of genes in the accessory genome for the different COG categories was measured with a chi-square correlation.

To document the overall taxonomic diversity of our genome collection, we built genus-level phylogenies for the core species (Fig. 3B/C/D). Firstly, we observed that the isolates and MAGs cluster together, confirming that we have isolated strains matching those identified within the kefir communities. Moreover, the genomes of *L. ghanensis* clustered into two clades, corroborating the presence of two strains previously described in Fig. 2C/D.

To identify gene content variation between strains we compared the genomes belonging to the same species in our genome collection. Overall, we annotated between 2041 and 2983 genes per species, of which 20-49% varied between genomes (i.e. accessory gene content) depending on the species (Fig. 3E). The fraction of shared genes rapidly decreased the more dissimilar the genomes were, independent of the completeness of the genome assembly (Fig. 3F).

To understand the functional significance of the accessory genomes, we tested which Clusters of Orthologous Genes (COG) categories were enriched in the accessory fraction compared to the core genome. Key cellular processes such as translation and nucleotide metabolism were very rarely observed in the accessory genome (Fig. 3G). On the contrary, the two functional groups significantly enriched (Chi-square test, p-value<0.01) in the accessory genomes of different core species were carbohydrate metabolism/transport and defense mechanisms (Fig. 3G). The variation of defense mechanism genes may be due to sample-specific phage infections, whereas the variation in carbohydrate metabolism and transport genes may be linked to different nitrogen sources used in the experiment. However, neither the Cazymes nor the defense mechanisms were significantly enriched in the genomes originating from specific inocula or nitrogen sources (Chi-square test, p-value>0.05, Supplement Fig. 10).

In summary, we found that closely related strains of core species of the analyzed water kefir samples show large variation in accessory gene content including functionally relevant gene categories (sugar utilization pathways and defense systems), but also that these were not associated with distinct treatment groups.

### Highly variable plasmids are present across water kefir samples and enriched in phage defense systems

Notably, we identified a total of 74 plasmids across the 14 isolate genomes, with *L. nagelii* and *L. paracasei* having on average six co-occuring plasmids within one isolate (Fig. 4A). In the metagenomes we assembled 409 high-quality (geNomad score>0.8) plasmids (de-replicated at >90% ANI). For many of them, we were able to assign the host species by an hierarchical classification tool named HOTSPOT (Ji et al. 2023). Most plasmids were predicted to be present in *Acetobacter s*pp. (n=105) and the fewest in *Liquorilactobacillus* spp. (n=8) (Fig. 4B). We found on average 88 plasmids per metagenomic sample with no significant differences between the different inocula or nutrient treatments (ANOVA, p-value=0.3, Fig. 4C). A total of 9% of the plasmids were found in three genera encompassing the core species (Supplement Fig. 11). Their prevalence showed a trimodal distribution with, on average, 34% of the plasmids being rare (<25% of samples), 34% occurring occasionally (25-75% of samples) and 32% were core (>75% of samples) (Fig. 4D).

**Figure 4.**
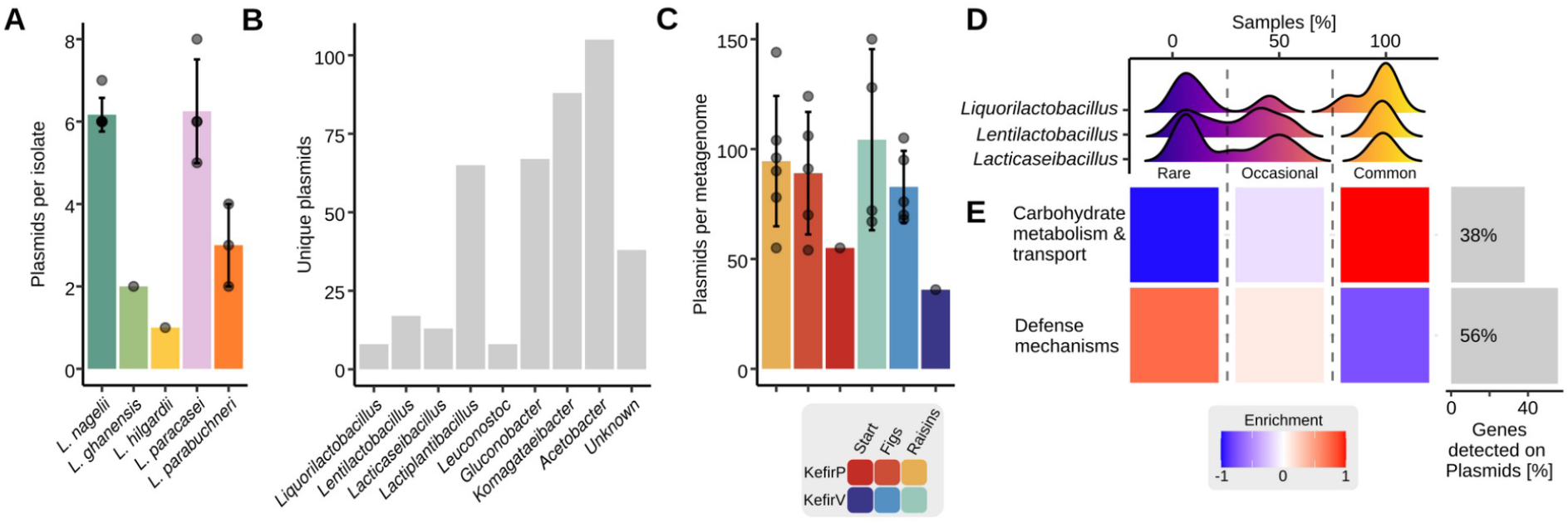
The distribution and functions of the plasmids within the water kefir samples. A) The number of plasmids detected in the different isolate genomes (grouped by species). B) The number of unique plasmids that are associated with the previously identified genera. C) The number of plasmids per metagenome (grouped by treatment). D) The prevalence of the plasmids in the different samples for the core genera (grouped into rare, occasional, and common plasmids). E) The fraction and enrichment of functions that were detected in the plasmids.

Interestingly, 38% of the accessory genes annotated as carbohydrate metabolism & transport and 56% of the genes annotated as defense mechanisms were located on plasmids (Fig. 4E). Moreover, carbohydrate metabolism & transport genes were primarily enriched (χ2 test, p-values<0.05) in the common plasmids, whereas the defense mechanisms were enriched in the rare plasmids (Fig. 4E).

### Sample-specific phage diversity

Phages can modulate the function and diversity of microbial communities (Koskella and Brockhurst 2014; Castledine and Buckling 2024). As we observed variation in gene content related to phage defense systems across genomes and samples, we hypothesized that active phages, potentially sample-specific phages, are present in water kefir communities. To identify and annotate phages from our samples, we analyzed all metagenomic contigs (> 4 kb) using the phage detection tool geNomad. We identified 203 dereplicated (at 95% sequence identity) phages over the 22 metagenomic samples (Fig. 5A). A total of 162 of these phages were predicted to have a lytic lifestyle, while 41 were identified as temperate phages. The phages showed a high level of variation in both G+C content (ranging from 16% to 66%) and in length (ranging from 4 kb to 141 kb, Fig. 5A). Every metagenomic sample contained on average 35 different phages (ranging from 18 to 55, Supplement Fig. 11). Approximately half (51%, 103/203 phages) of the identified phages were found in two or more samples (Fig. 5B), whereas three temperate phages were identified in every sample. Interestingly, 100 phages (49%) were sample-specific. While two samples had no sample-specific phages, many had several with up to 13 unique phages in sample K30 (Fig. 5B inset plot). There was no apparent variation according to inocula nor nutrient treatment in the number or type of phages detected across samples (Fig. 5B).

**Figure 5.**
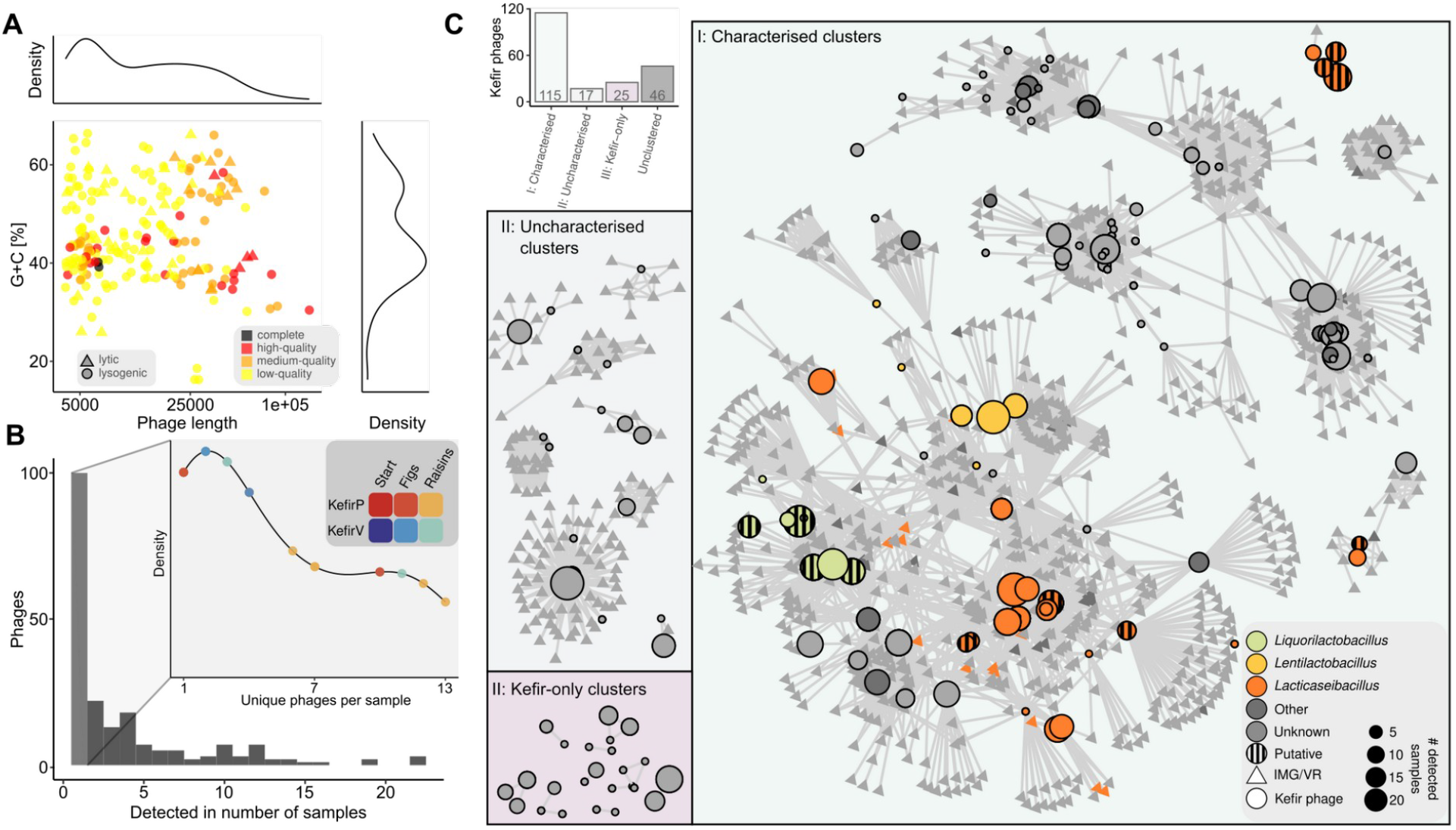
The diversity, distribution, and host classification of phages in water kefir. A) The diversity of the dereplicate phage assemblies from all water kefir metagenomes. Additionally, we show the completeness and lifestyle of the phages. The density plots show the distribution of the data along the axes. B) Phage distribution, shown as the number of metagenome samples in which each dereplicated phage was identified. The inset plot highlights sample-specific phages and the samples in which they occur. C) The clustering of all dereplicated phages and related IMG/VR phages. The phage host assignments and the number of samples that the phage is detected in are illustrated accordingly. Moreover, the number of phages in the different categories, namely I: characterised clusters, II: uncharacterised clusters, III: kefir-only clusters or unclustered, is illustrated in the accompanying barplot in the top left.

Next, we wanted to know the bacterial host of these phages. To do so, we created an all-against-all gene clustering using Vcontact2 (Bin Jang et al. 2019). We clustered the assembled phages from our water kefir samples in combination with the extensive IMG/VR phage reference database (Camargo et al. 2023). In total, 77% (157/203) of all phages detected in the water kefir samples fell into a cluster. Only 46 phage contigs remained singletons (unclustered) (Fig. 5C). A total of 115 clustered phages included one previously characterised phage, 17 were uncharacterized but included phages from the reference database, and 25 clusters consisted of only phages from our water kefir samples. Using the tool iPHoP (Roux et al. 2023), we then assigned a putative bacterial host to each characterized phage. A total of two, four, and sixteen phages were predicted to infect bacteria of the three core genera *Liquorlactobacillus, Lentilactobacillus*, and *Lacticaseibacillus*, respectively. An additional 16 phages were also predicted to infect these hosts, because they clustered with previously identified phages targeting these hosts (striped circles in Fig. C). Moreover, we identified numerous phages targeting genera considered to be present occasionally or rarely across our samples: *Lactobacillus* (n=16), *Acetobacter* (n=10), *Leuconstoc* (n=3) and *Streptococcus* (n=2) (Supplement Fig. 12). Interestingly, we also identified a substantial number of phages potentially infecting bacterial genera that were not detected in our 16S rRNA gene sequencing analysis (Supplement Fig. 12) but that have been commonly identified in water kefir samples in other studies, namely *Oneoccocus* (n=10), *Ligilactobacillus* (n=11), and *Limosilactobacillus* (n=11) (labelled “other” in Fig. 5C). We hypothesize that these phages either have a broader host range than previously expected (Hwang et al. 2023) or that these genera were present but not detected by our analyses.

## Discussion

This study provides novel insights into the diversity and dynamics of microbial communities beyond the species-level in water kefir. We demonstrated that i) species-level stability is maintained over short and long timescales, ii) SNP diversity within and between samples is generally low, iii) gene content variation is driven by accessory genomes, particularly plasmids, and iv) diversity might be influenced by phage-driven dynamics. By highlighting the interplay between plasmids, phages, and accessory gene content, our work underscores the importance of studying microbial diversity beyond the species-level to understand the ecological and evolutionary processes shaping microbial communities.

We observed remarkable yeast and bacterial stability at the species-level, with only a few accessory species replaced between two water kefir starter cultures that were split about two years apart. Firstly, among yeasts, we consistently detected the two species commonly identified in water kefir, *Saccharomyces cerevisiae* and *Brettanomyces bruxellensis* (Laureys *and De Vuyst 2014). These species are known for their roles in flavor-related processes, with B. bruxellensis*, in particular, contributing to a pungent flavor profile through its metabolic activity (Steensels and Verstrepen 2014; Crauwels et al. 2015). A similar trend was observed for bacteria. The most abundant species were consistently present in all samples and corresponded to community members described to be prevalent in previous studies (Fels et al. 2018; Magalhães et al. 2010; da C. P. Miguel et al. 2011; Laureys et al. 2018; Verce, De Vuyst, and Weckx 2019; Laureys and De Vuyst 2014). A greater variability was found among some of the rare species such as *Bifidobacterium aquikefiri* and *Lactobacillus parabuchneri* which were restricted to one inoculum. These species are particularly interesting due to their respective associations with putative probiotic activity (Verce, De Vuyst, and Weckx 2019) and flavor compound production (i.e histamine) (de A Møller et al. 2021). Notably, several bacterial species reported to be commonly present in water kefir were absent from our samples, such as *Zymomonas* spp. and *Oenococcus spp. (Marsh et al. 2013; Verce, De Vuyst, and Weckx 2019; Patel et al. 2022)*. Instead, we found more acetic acid bacteria in our samples. The prevalent species exhibited slightly more variability in their absolute abundances in KefirV samples compared to KefirP over a short timescale, indicating that KefirP may be more reproducible. Moreover, in two cases we saw a substantial shift with more acetic acid bacteria identified. Water kefir fermentation consists of a microbial succession, where acetic acid bacteria are often late arrivals, profiting from the metabolites secreted by yeasts and lactic acid bacteria (Pendón et al. 2022). Therefore, these differences may be explained by minor changes in the fermentation time across replicates (Arrieta-Echeverri et al. 2023; Martínez-Torres et al. 2017).

Strain-level diversity within samples was generally low, consistent with previous observations in other fermented foods such as cheese starter cultures (Somerville et al. 2024), but varied significantly across the core bacterial species. Since these species were all abundant and core community members, population size alone does not explain the observed differences in strain-level diversity. A more likely explanation is the variability in ecological niches that each bacterial species can occupy. Among the five core species, *Lentilactobacillus hilgardii* exhibited the lowest SNP diversity. This species is known for its role in exopolysaccharide production, a key factor in maintaining the structural integrity of kefir grains (Waldherr et al. 2010). In this case, the low strain-level diversity could indicate that only a specialized subpopulation is capable of efficiently producing this compound. Alternatively, the persistent and close co-existence within kefir grains may foster interdependencies between specific strains, thereby restricting strain-level diversity, as has been observed in milk kefir (Blasche et al. 2021). Interestingly, KefirP samples, while being more stable at the species level, displayed significantly higher strain-level diversity compared to KefirV, with SNP frequencies being 10 to 1,000 times higher. Higher strain-level diversity could potentially facilitate adaptation to different ecological conditions (Gibbons and Gilbert 2015) and could potentially indicate that this kefir has previously been grown on more variable nitrogen sources. When comparing strain-level diversity between samples, only over the long timescale, strain replacement was observed in two of the five species, highlighting the resilience of these microbial communities to strain invasion. While cultivation has often been applied to identify species diversity in water kefir samples (Zannini et al. 2022), only the collective analysis of isolate culturing (14 isolates) and in-depth shotgun metagenomic strain analysis has enabled us to shed light on strain-level diversity. Despite this low nucleotide diversity, the accessory genome was enriched with genes related to sugar utilization and phage defense, many of which were encoded on plasmids. On average, the isolate genomes of the core species contained six co-existing plasmids. This is consistent with previous findings in other fermentation species such as *Lactococcus lactis*, where large numbers of plasmids facilitate rapid adaptation to changing environments (van Mastrigt et al. 2018). While we do not know much about plasmid diversity and dynamics in water kefir, the high variability in plasmid content highlights their potential role in driving strain-specific adaptations, particularly in response to phage pressure or shifts in substrate availability. Notably, phage defense mechanisms have been shown to vary rapidly even among closely related strains (Hussain et al. 2021) also in dairy fermented foods (Somerville, Schowing, et al. 2022). Future research could focus on the potential importance of plasmids as dynamic genetic elements driving the resilience and adaptability of microbial communities in fermented systems.

While previous studies have highlighted the variability of different kefir samples (Hsieh et al., 2012; Marsh et al., 2013; da Miguel et al., 2011), we have seen remarkable stability when following the same water kefir in response to a change in nitrogen source. Microbial changes rarely happened by species or strain replacement but rather by sweeps of genomic elements, notably plasmids. To fully understand the selection pressure and dynamics shaping specific gene functions, future studies should track microbial communities with higher taxonomic (beyond dominant strains), temporal (beyond endpoint analysis) and spatial resolutions. The granular structure of water kefir grains likely plays a key role in this stability. The microbes usually grow on the outside of the grains (Neve and Heller 2002). This structure could limit extensive plasmid mobilization by creating microenvironments around individual cells. Additionally, the grain’s physical structure may enhance resilience against microbial and phage invasion, as previously observed in biofilms (Kunisch et al. 2024).

Phages appear to play a pivotal role in shaping the bacterial accessory genome. We identified an average of 35 distinct phages per sample, with nearly half being sample-specific. Sample-specific phage infections indicate that phages may be introduced from different culturing environments (student households) rather than systematic differences between nitrogen sources. This high phage diversity suggests dynamic interactions between phages and bacterial hosts, likely influencing the distribution of phage defense mechanisms within the community. Moreover, by demonstrating that metagenomic approaches can identify phages, both isolated and non-isolated phage clusters targeting the different bacterial host taxa, this study provides a strong foundation for exploring the diversity, dynamics, and roles of phages in traditional food fermentation systems. Overall, this remarkable stability underscores the resilience of water kefir communities to environmental perturbations as well as microbial and viral invasion, both over short ecological timescales and largely over longer evolutionary timescales.

Understanding microbial diversity beyond the species-level is key to unraveling the dynamics and evolution of microbial communities (Van Rossum et al. 2020). Using water kefir as a model, we explored microbial diversity changes over long and short timescales, uncovering insights into species- and strain-level diversity, gene content variability, and the role of plasmids and phages. Future studies focusing on plasmid content and dynamics will provide insights on the role of plasmids towards the rapid adaptation to environmental changes in nutrients and phages.

In addition to advancing our scientific understanding, this study also served as a powerful educational experience for microbiology students. Through hands-on training in state-of-the-art bioinformatic techniques, including 16S rRNA sequencing, shotgun metagenomics, and genome assembly, students not only contributed to meaningful research but also gained a deeper appreciation for the complexities of microbial ecosystems. Leveraging accessible and relatable systems like food fermentation to inspire and educate the next generation of scientists is vital for fostering curiosity, creativity, and innovation in the biological sciences. This work demonstrates that, through collaboration and education, we can push the boundaries of knowledge while empowering future researchers to tackle the many challenges of microbial ecology and evolution.

## Methods

### Kefir passaging

The two water kefir inocula (KefirP and KefirV) were distributed in falcon tubes to 35 graduate students. Further, the students received 250g of standardized sugar and an organic lemon. Finally, each student was randomly assigned to receive organic figs or raisins. The students were instructed to do five passages, each 36 h containing 0.5 L of tap water, 30g/L sugar, one lemon slice, and either half a fig or 25 raisins. This was inoculated with two tablespoons of water kefir grains. The students collected the five supernatants, marked the grain growth and after the five passages collected the grains (see details in *“Supplement method I: Passaging protocol”*).

### Strain isolation, identification and DNA extraction

Kefir grains from the initial kefir culture (KefirV) were broken down in a Stomacher blender (Interscience, USA) and plated on MRS, ML, and M17 plates. The plates were incubated at 30 °C for 3 and 8 days. Colonies with different morphologies were picked, re-plated, eventually grown in liquid media and stored in 20% glycerol at −80°C (see details in *“Supplement method II: Colony picks”*). The species identification of the selected colonies was performed on the MALDI-TOF as previously described (Lüdin et al. 2018). The DNA of the colonies was extracted with the EZ1 DNA Tissue kit on the BioRobot EZ1 robot (Qiagen, Hombrechtikon, Switzerland).

### Water kefir DNA extraction

Water kefir grains were ground frozen to fine powder in a 50 ml grinding jar with a 25 mm ball (stainless steel) in a Mixer Mill MM 400 (Retsch, Switzerland), before DNA isolation. Thereafter, crude DNA was extracted using custom protocol and further purified either with the Zymoclean Large Fragment DNA recovery kit (Zymo Research) or DNeasy PowerSoil kit (Qiagen) and CleanNGS magnetic beads (Clean NA). For step-by-step protocol see *“Supplement method III: DNA extraction protocol”*

### Genome and metagenome sequencing

Both the isolate genomes and the water kefir metagenomes were sequenced on a Illumina Novaseq. Nextera flex libraries were prepared and subjected to Novaseq 150PE (Illumina) sequencing at the Genomic Technologies Facility of the University of Lausanne (Switzerland) and at Novogene (Cambridge, UK). Further, we sequenced the genomes with the rapid barcoding kit on a minION (Nanopore) (R9.4) at the IFIK (Bern, Switzerland).

### 16S community analysis

Region V4 of the 16S rRNA gene was amplified using the primers 515F-Nex and 806R-Nex (Caporaso et al. 2011) as described previously (Kešnerová et al. 2017). PCR product concentrations were quantified by PicoGreen and pooled in equimolar concentrations to sequence on an Illumina MiSeq sequencer (2 × 250 bp) by the Genomic Technology Facility of the University of Lausanne. The statistical analysis of the data was conducted with the DADA2 pipeline (see details in script) (Callahan et al. 2016). Alpha- and beta-diversity was calculated with phyloseq (McMurdie and Holmes 2013) and vegan R packages (Dixon 2003).

### Genomic and metagenomic raw read analysis and assembly

All Illumina raw reads were quality- and adaptor-trimmed with cutadapt (Martin 2011). Mapping to the reference database and identification of human contamination was done with bwa mem (H. Li 2013). SNPs were called with freebayes-parallel (Garrison and Marth 2012). The genomes and metagenomes were assembled with SPAdes (Bankevich et al. 2012) and Flye (Kolmogorov et al. 2019). The Flye assemblies were polished with four rounds of Racon (Vaser et al. 2017) polishing and four rounds of freebayes polishing (Garrison and Marth 2012). The completeness of the metagenome assemblies was checked with Quast (Mikheenko et al. 2023) and checkM (Parks et al. 2015). The assemblies were submitted to NCBI (see data availability section). All against all mapping against the assemblies was done with bwa mem (H. Li 2013). The read coverage was calculated with jgi_summarize_bam_contig_depths and binning was done with metabat2 (Kang et al. 2019) MAG quality and taxonomy was inferred by CheckM (Parks et al. 2015) and GTDBTK (Chaumeil et al. 2019).

### Metagenomic taxon profiling and SNP analysis

Metagenomic taxon profiling for kingdom, yeast species and *Liquorilactobacillus* species diversity was conducted with metaphlan (Truong et al. 2015) and kraken2 (Wood, Lu, and Langmead 2019). The relative abundances of the three core *Liquorilactobacillus* species were used to further distinguish the different species from the 16S rRNA sequencing data. To do strain-level diversity analysis we map against a reference database. The reference database consisted of sequenced isolate genomes from our kefir or NCBI strains (see Zenodo supplement). Freebayes (Garrison and Marth 2012) and SNPeffect (Baets et al. 2012) were used to identify core synonymous/non-synonymous mutations. Distances between two samples were measured with Bray-Curtis distance and illustrated with a rTSNE (Maaten 2014). PERMANOVAs were generally calculated with the adonis2 function from the r vegan package (Dixon 2003). Additionally, inStrain was used to estimate average nucleotide identity (estANI) (Olm et al. 2021).

### Gene content analysis and phylogenies

Gene annotation of bacterial chromosomes and plasmids was done with BAKTA (Schwengers et al. 2021), while phage genomes were annotated with Pharokka (George et al. 2023). Additionally, the gene categories (e.g. COG) were annotated with the EGGNOG database (Rodríguez Del Río et al. 2023). The defense mechanisms were annotated with DefenseFinder (Tesson et al. 2022). Pangenome analysis and core proteome phylogenies were created with Orthofinder (Emms and Kelly 2019). Average nucleotide identities were calculated with fastANI (Jain et al. 2018). Gene enrichment analysis was done by chi-square (r-base) test of the prevalence of different CAZymes and phage defense genes in the MAGs from the two nitrogen sources.

### Plasmid and phage analysis

Phages and plasmids were identified with geNomad (Camargo et al., n.d.). Only phages and plasmids with quality scores above 0.8 and longer than 4 kb were kept. The contigs were dereplicated with cd-hit est and 95% identity and 85 % coverage cutoff (W. Li and Godzik 2006). The phage completeness was evaluated with checkV (Nayfach et al. 2021). Plasmid hosts were identified with hotspot (Ji et al. 2023). Phage hosts were inferred by clustering together with the IMG VR database using vcontact2 (Bin Jang et al. 2019) and transferring of all cluster information.

### Statistics, scripts and data

All statistics were done within R (R Core Team, 2020) and ggplot2 (Wilkinson 2011). All corresponding code for the bioinformatics, statistics and plotting is available on github (https://github.com/Freevini/2025_Kefir). The data is deposited on zenodo (10.5281/zenodo.14940280) and NCBI (genomes bioproject PRJNA1203286 and metagenome bioproject PRJNA1229503).

### Lectures and Tutorials

This manuscript was created based on data, material and analysis prepared for a graduate course at the University of Lausanne in 202120/22 and 2022/2023. All course resources have been made available on github (https://github.com/Freevini/SAGE_21-23).

## Supporting information

Supplement figures_ Waterkefir

Supplement method I_ Passaging protocol

Supplement method II_ Colony picks

Supplement method III_ DNA extraction protocol

## Acknowledgments

We thank all the SAGE course students from 21/22: Alexandre Jann, Alexis Dentand, Antonio Garrido Marques, Béatrice Pichon, Benjamin Skaggs, Camille Schmidt, Cyril Ben Amara, Daniel Hafez, Elindi de Coning, Estelle Juttin, Estelle Vivien, Giorgia Wennubst Pedrini, Jeronimo Camelo, Julien Cergneux, Karunnya Tharmakulasinkam, Laetitia Holzer, Léonie Mottet, Margaux Corset, Mathilde Buvelot, Mohamed Abdo, Nicolò Tartini, Omar Keshk, Petros Liakopoulos, Philip Gwyther, Samuel Moix, Sarah Abou-zite, Xiaojing Peng and SAGE 22/23 : Abhay Singh Alex Taimsalu, Aline Altenried, Andrea Perugini, Antoine Induni, Arun Maurya, Delphine Studer, Derek Scott, Giulia Saitta, Jana Brenner, Jérôme Benedetti, Juan Rueda, Léo Moser, Loïc Zen-Ruffinen, Manon Schmidli, Sarah Moreira Clemente, Théo Schneider, Wan-Ting Huang, Xavier Vuattoux. Moreover, we thank Noam Shani and Ueli vonAh for helping in the initial Waterkefir strain isolation. Also we thank the Lausanne Genomic Technologies Facility (GTF) for the continuous help with sequencing. The authors would like to thank the SNF Postdoc.Mobility P500PB_214419 (V.S.). This work was supported as a part of NCCR Microbiomes, a National Centre of Competence in Research, funded by the Swiss National Science Foundation (grant number 180575).

## Supplement

- Supplement figures:
- Supplement method I: Passaging protocol
- Supplement method II: Colony picks
- Supplement method III: DNA extraction protocol

