## Supplement figures_ Waterkefir for "Strain-level and phenotypic stability contrasts with plasmid and phage variability in water kefir communities"

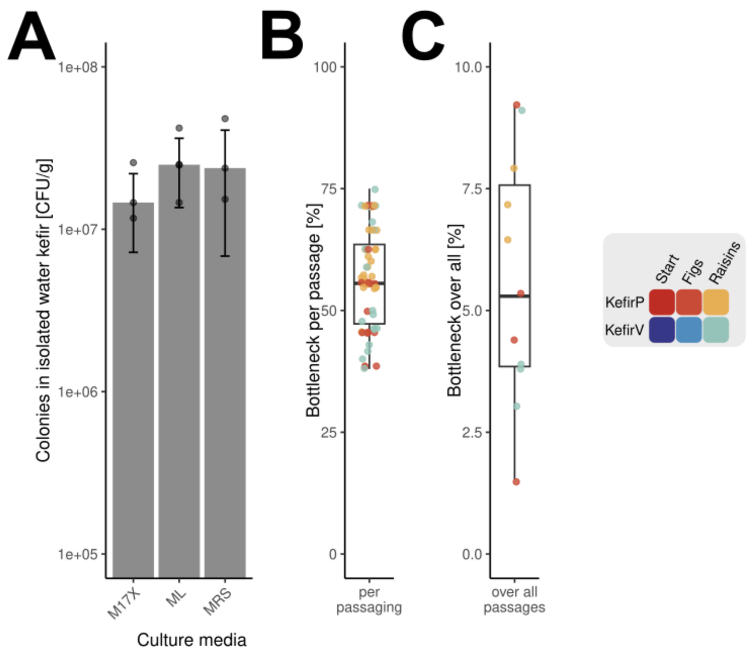


Supplement figure 1. A) The colony forming units (CFU) of the water kefir on three different growth media. B) The average bottleneck per passage calculated for the water kefir experiment. C) The overall bottleneck calculated for the water kefir experiment. The color of the dots indicates the treatment group (see legend).

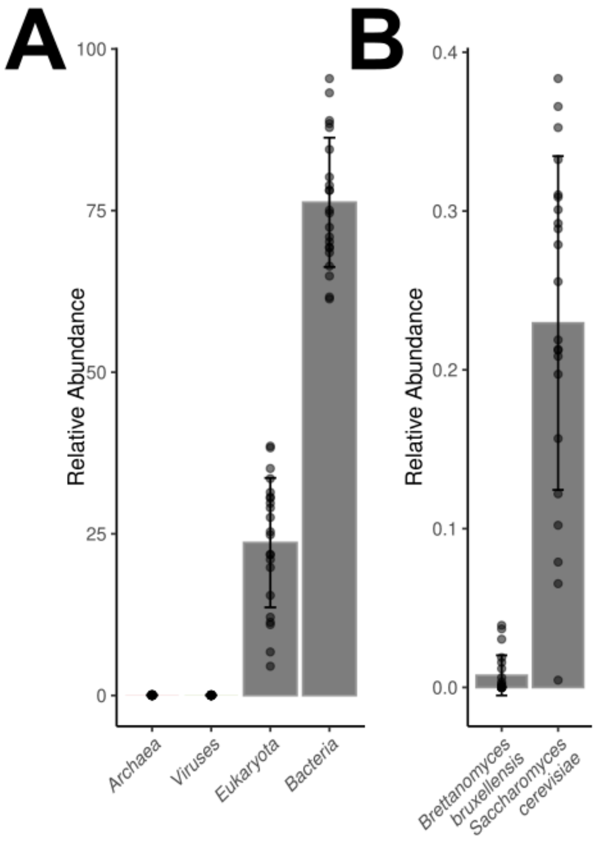

Supplement Figure 2. A) Microbial diversity in the different metaG samples on the kingdom level. B) The eukaryotic microbial diversity of the two yeast species.

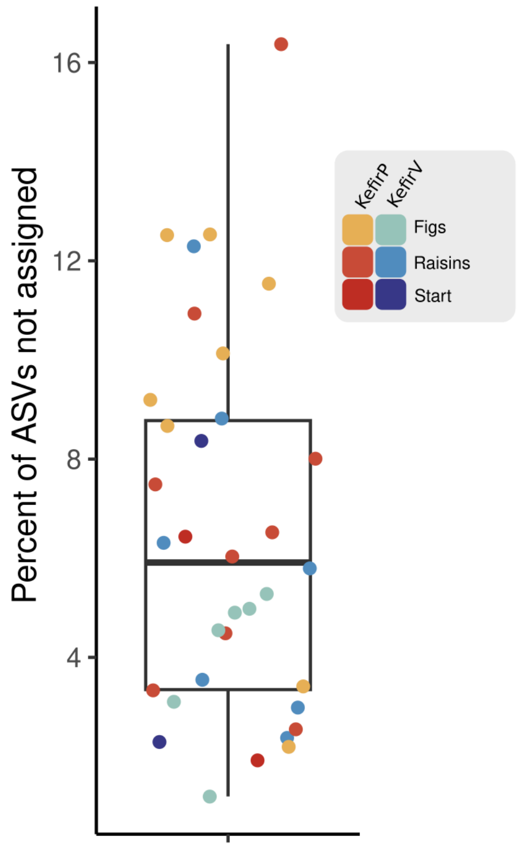


Supplement Figure 3. The boxplot of the percent of ASVs that are either not assigned to a genus or are below 0.1% relative abundance. The color of the dots indicates the treatment group (see legend).

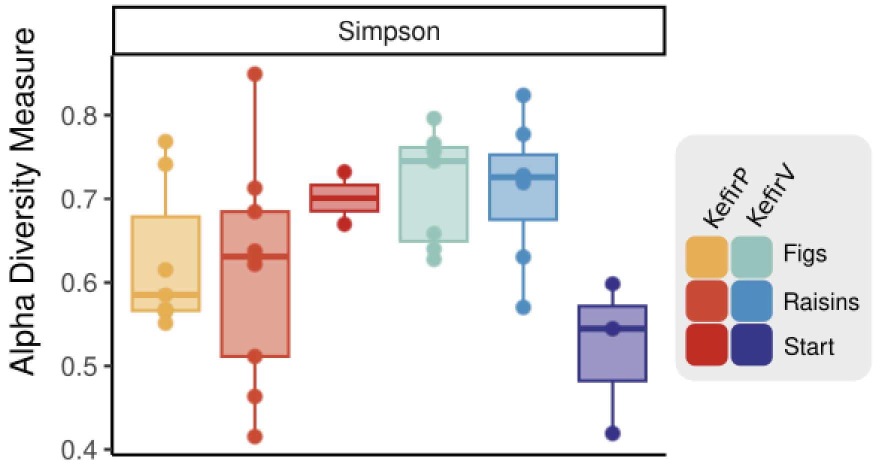


Supplement Figure 4. Alpha diversity was measured with the Simpson index for the different 16S rRNA samples. The columns indicate the different treatment groups.

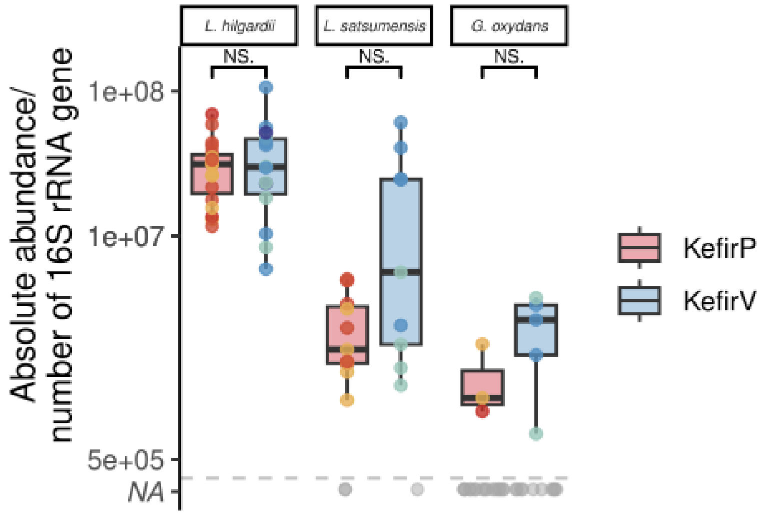


Supplement Figure 5. Absolute abundance of bacterial species with similar abundances in KefirV and KefirP samples. Each dot represents a different student sample (biological replicate). Statistical significance is indicated above the boxplots (NS = not significant; *** = p-value < 0.01).

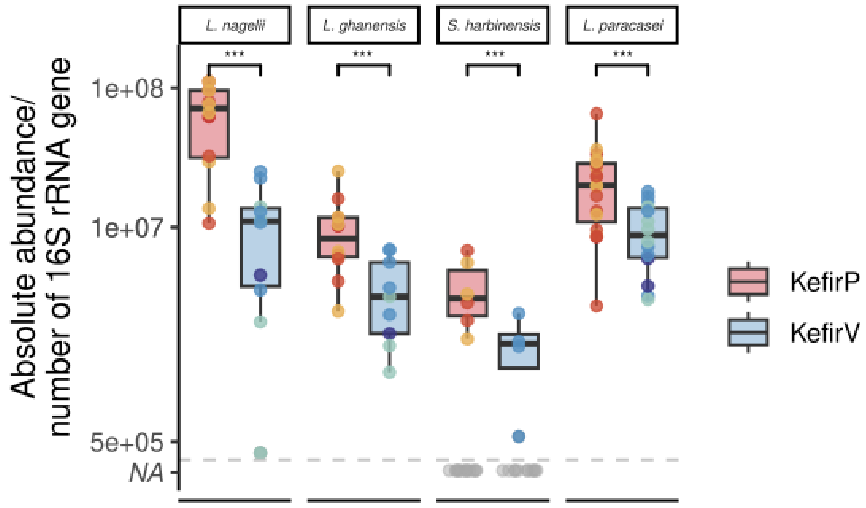


Supplement Figure 6. Absolute abundance of the bacterial species that are more abundant in the KefirP samples. Each dot represents a different student sample (biological replicate). Statistical significance is indicated above the boxplots (NS = not significant; *** = p-value < 0.01).

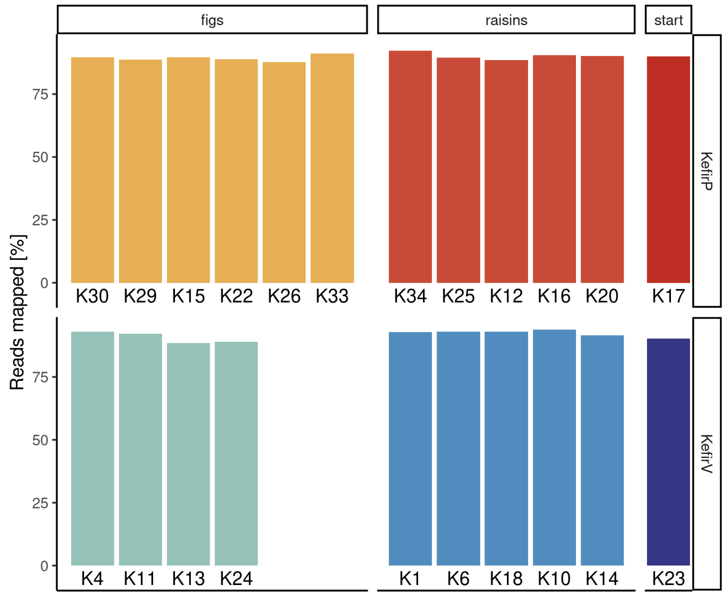


Supplement Figure 7. The percent of reads mapped to the complete reference genome collection.

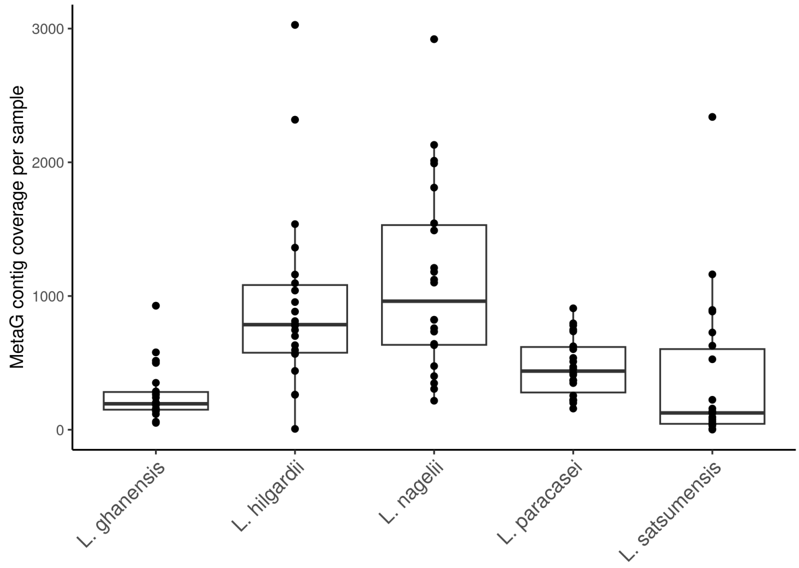


Supplement Figure 8. Contig coverage for the different samples and core species in the metagenomic samples.

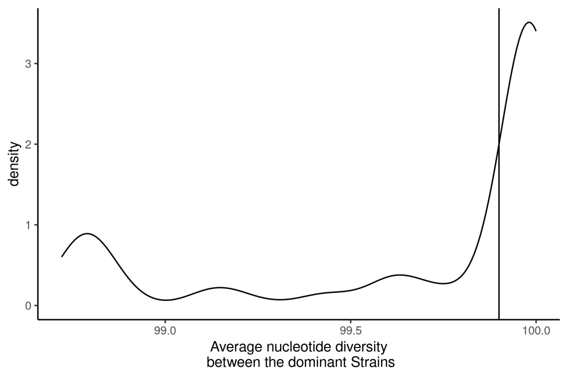


Supplement Figure 9. The overall distribution of the average nucleotide diversity within the core species. The vertical line indicates the 99.9% threshold.

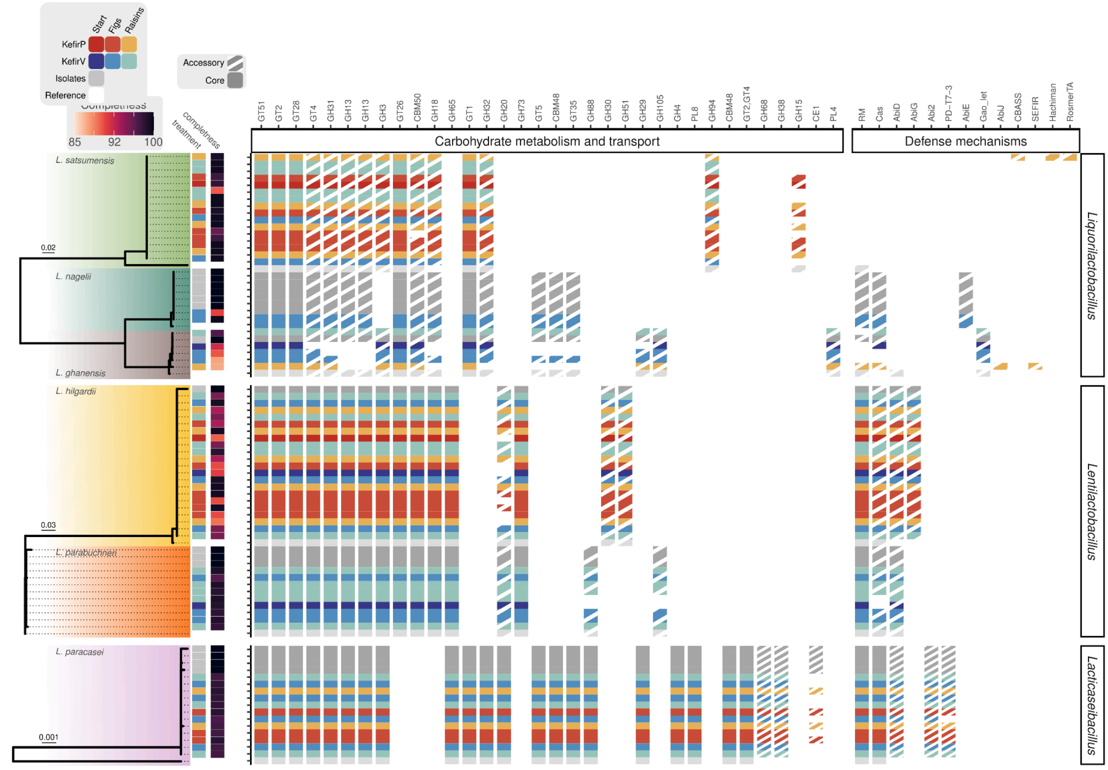


Supplement Figure 10. The genome phylogenies of the three core genera, including all isolate genomes and MAGs and one reference strain per species. The completeness and which treatment the MAG originates from are illustrated in the adjacent heatmap. The presence and absence of the different carbohydrate metabolism & transport (Cazyms) and ii) defense COGs for the different strains in the genome collection. The colours correspond to the adjacent phylogeny and the pattern corresponds to the core and accessory patterns (see legend top left).

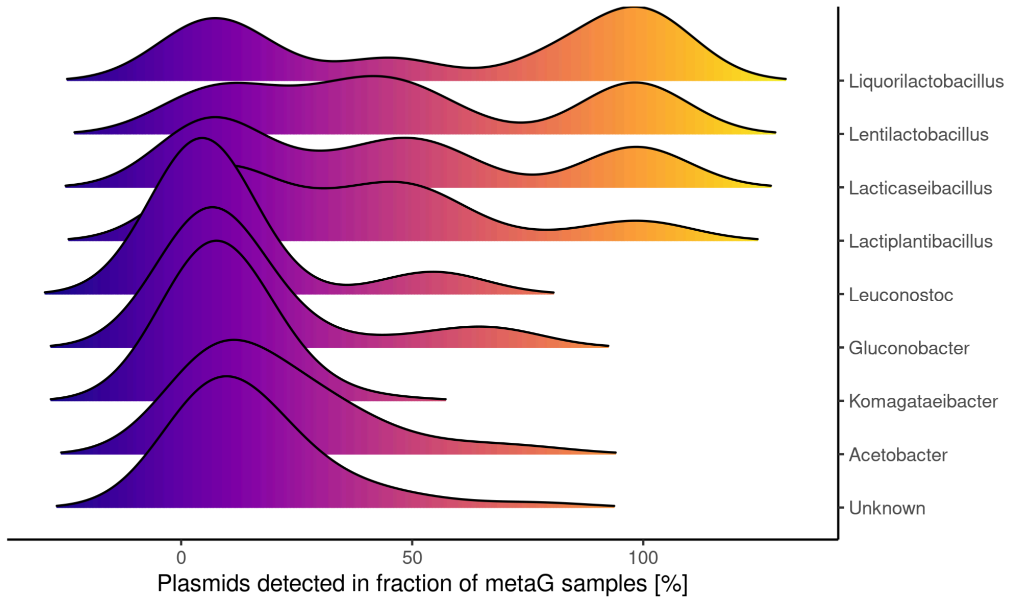


Supplement Figure 11. The fraction of metagenomic samples in which plasmids associated to the different genera were detected in.

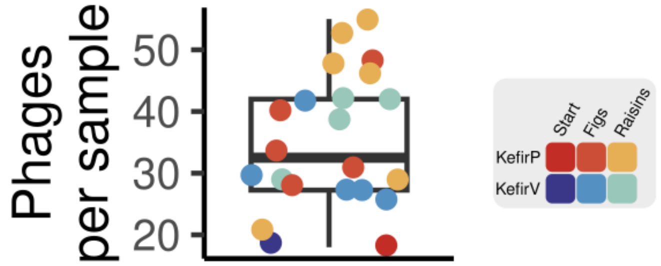


Supplement Figure 12. The number of phages observed in the different samples with the coloring of the points according to the sample it came from. There is no significant difference between the treatments (Wilcoxon test P-value > 0.05).

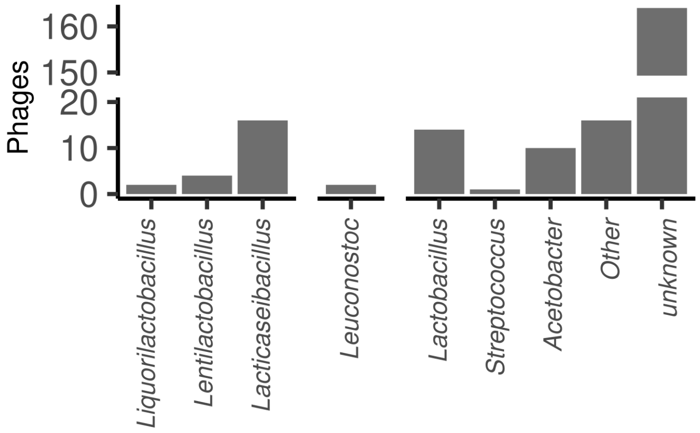


Supplement Figure 13. Phage host prediction from iPHoP for the different core, occasional and rare genera. Moreover, the category “other” includes genera not identified with 16S.
