## Supplement method I_ Passaging protocol for "Strain-level and phenotypic stability contrasts with plasmid and phage variability in water kefir communities"

### Basic Recipe

- 30 g of sucrose
- 500 ml of tap water
- 25 raisins or 1/2 fig
- 1 slice of lemon
- 2 tablespoons of water kefir grains

### Basic workflow

**1.Start kefir brewing on day 1**

- Fill sugar to the line indicated on the plastic cup and add to fermentation glass jar (this equals 30 g)
- Fill 500 ml of tap water into the jar (if you don't have a measuring cut you can measure and label 8 cm from the bottom)
- Dissolve sugar with spoon
- Add 25 raisins and the lemon slice
- Add kefir grains using the same line as for the sugar indicated on the plastic cup
- Let it incubate at room temperature (i.e. ferment) for 36h (e.g. start at 8 am in the evening and stop at 8 pm the following day)

**2. Stop primary fermentation on day 2**

- Remove figs and lemon with the strainer
- Fill up the Eppendorf tube with the water kefir liquid and put it in the freezer (labelled: NAMEnum_passageNUMBER)
- Collect the kefir grains with the strainer, wash with tap water and put into a plastic cup to measure the volume of  grains (indicated on the cup)
- Collect kefir grains in a separate glass jar by adding twice as much volume of tap water, and store in the fridge, or start another kefir fermentation using the same volume as on Day 1 (kefir grains reproduce during fermentation, keep the leftovers or trash)

**Optional in case you want to consume your kefir lemonade**

- Use strainer + funnel to fill liquid without grains into a 0.5 L PET bottle
- Close PET bottle tightly (bottle can be squished so no air is in anymore). Now you let it ferment for one more day (24).
- Then put it into the fridge before drinking it.
