## Supplement method II_ Colony picks for "Strain-level and phenotypic stability contrasts with plasmid and phage variability in water kefir communities"

| colony | time after plattir | media | size | info | kept | Maldispecies | DNA/sequence | expected genon | Vazyme DNA co | HMW DNA Conc | Send to GTF | ESL number | metaphlan | contamination | spades assemb | spades assemb | spades assembly contigs | *blasting the longest contigs* | closest fastANI | ANI | coverage_ANI | not sent? | ONT_first_roun | ONT_longest co | ONT_contigs_n |
|---|---|---|---|---|---|---|---|---|---|---|---|---|---|---|---|---|---|---|---|---|---|---|---|---|---|
| 5 | 3days | MRS | big | whitish | yes | *L. paracasei* | | | 16 | 31.4 | yes | ESL0961 | Lactobacillus_casei_paracasei | | 3106144 | | 230 | *Lacticaseibacillus paracasei* | ESL0969 | 99.9889 | 0.990788 | | Y | 3057602 | 6 |
| 7 | 3days | MRS | small | | yes | *L. parabucherni* | yes | | 4.88 | 44.9 | yes | ESL0962 | unclassified | | 2572293 | | 72 | *Lentilactobacillus parabuchneri* | ESL0970 | 99.9993 | 0.997608 | x | N | | |
| 8 | 3days | MRS | small | | yes | *Lactobacillus gha* | yes | | 43.7 | 50.6 | yes | ESL0963 | unclassified | | 2432514 | | 53 | *Latilactobacillus curvatus/plantarun* | ESL0974 | 80.1245 | 0.563291 | | Y | 2468184 | 2 |
| 9 | 3days | ML | big | white | yes | *L. nagelli* | Yes | | 27.9 | 34.9 | yes | ESL0964 | unclassified | | 2425112 | | 72 | *Liquorilactobacillus nagelii* | ESL0974 | 99.9395 | 0.970738 | | Y | 2336742 | 5 |
| 11 | 3days | ML | big | transparent | yes | *L. paracasei* | | | 10.8 | 13.4 | yes | ESL0965 | Lactobacillus_casei_paracasei | | 3093839 | | 187 | *Lacticaseibacillus paracasei* | ESL0969 | 99.9916 | 0.985626 | x | N | | |
| 13 | 3days | ML | small | white | yes | *L. nagelli* | yes | | 14.4 | 20.2 | yes | ESL0966 | unclassified | | 2442172 | | 68 | *Liquorilactobacillus nagelii* | ESL0974 | 99.9918 | 0.994949 | x | N | | |
| 15 | 3days | ML | small | transparent | yes | *L. paracasei* | | | 9.37 | 14 | yes | ESL0967 | Lactobacillus_casei_paracasei | | 3097836 | | 177 | *Lacticaseibacillus paracasei* | ESL0969 | 99.9685 | 0.988798 | | N | | |
| 16 | 3days | ML | small | transparent | yes | *L. parabucherni* | yes | | 4.84 | 12.6 | yes | ESL0968 | unclassified | | 2642694 | | 479 | *Liquorilactobacillus nagelii* | ESL0974 | 99.969 | 0.997478 | | N | | |
| 21 | 3days | M17 | big | white | yes | *L. paracasei* | | | 11.3 | 12.9 | yes | ESL0969 | Lactobacillus_casei_paracasei | | 3094339 | | 171 | *Lacticaseibacillus paracasei* | ESL0967 | 99.9922 | 0.990816 | | N | | |
| 26 | 3days | SC | big | high | yes | *L. nagelli* | | | 3.45 | 13.9 | yes | ESL0970 | unclassified | | 2673294 | | 84 | *Lentilactobacillus parabuchneri* | ESL0962 | 99.9976 | 0.963091 | | N | | |
| 27 | 3days | SC | big | high | yes | *L. paracasei* | yes | | 45.1 | 30.7 | yes | ESL0971 | unclassified | | 2453318 | | 56 | *Liquorilactobacillus nagelii* | ESL0974 | 99.9503 | 0.961298 | | Y | 2326109 | 5 |
| 28 | 3days | SC | small | | yes | *L. parabucherni* | yes | | 41.8 | 32.1 | yes | ESL0972 | unclassified | | 2416470 | | 54 | *Liquorilactobacillus nagelii* | ESL0974 | 99.9656 | 0.968274 | | Y | 2326005 | 6 |
| 29 | 3days | SC | small | | yes | *L. nagelli* | | | 1.28 | 14.8 | yes | ESL0973 | Saccharomyces_ | <0.5% Naumovo: | 13684538 | | 7940 | *Saccharomyces cerevisiae* | ESL0977 | 99.8955 | 0.947454 | | N | | |
| 30 | 3days | FH | small | | yes | *L. nagelli* | | | 57 | 47.3 | yes | ESL0974 | unclassified | | 2432891 | | 63 | *Liquorilactobacillus nagelii* | ESL0972 | 99.952 | 0.967172 | | Y | 2367332 | 6 |
| 35 | 8days | M17 | kleiner | | yes | Dekkera bruxelle | yes | | 0.575 | 9.1 | yes | ESL0975/ESL09! | Lactobacillus_casei_paracasei | | 16729850 | | 15123 | *Brettanomyces bruxellensis* | NA | NA | NA | | N | | |
| 47 | 8days | MRS | small | | yes | ***L. hildargii*** | yes | | 21.9 | 19.1 | yes | ESL0976/ESL0986 | | | 3252498 | | 145 | *Lentilactobacillus hilgardii* | ESL0970 | 78.5605 | 0.190058 | | N | | |
| 50 | 8days | MRS | big | | yes | ***L. hildargii*** | yes | | 10.4 | 39.8 | yes | ESL0977/ESL0989 | | | 12731775 | | 5110 | *Saccharomyces cerevisiae* | ESL0973 | 99.8618 | 0.904747 | | N | | |
| 1 | 3days | MRS | big | redish | No | ***no hit*** | | | | | Not FOUND | | | | | | | | | | | | | | |
| 2 | 3days | MRS | big | redish | yes | *S. cervisiae* | yes | | 7 | | | | | | | | | | | | | | | | |
| 3 | 3days | MRS | big | whitish | yes | *L. paracasei* | yes | | 14.3 | | | | | | | | | | | | | | | | |
| 4 | 3days | MRS | big | whitish | No | ***no hit*** | | | | | | | | | | | | | | | | | | | |
| 6 | 3days | MRS | small | | yes | *L. nagelli* | yes | | 120 | | | | | | | | | | | | | | | | |
| 10 | 3days | ML | big | white | yes | *L. nagelli* | | | 33.1 | | | | | | | | | | | | | | | | |
| 12 | 3days | ML | big | transparent | yes | *L. paracasei* | | | 0.65 | | | | | | | | | | | | | | | | |
| 14 | 3days | ML | klein | white | yes | *S. cervisiae* | | | | | | | | | | | | | | | | | | | |
| 17 | 3days | PY | big | | yes | *S. cervisiae* | | | | | | | | | | | | | | | | | | | |
| 19 | 3days | M17 | big | redish | yes | *S. cervisiae* | | | | | | | | | | | | | | | | | | | |
| 20 | 3days | M17 | big | redish | yes | *S. cervisiae* | | | | | | | | | | | | | | | | | | | |
| 22 | 3days | M17 | big | white | yes | *L. paracasei* | | | 2.71 | | | | | | | | | | | | | | | | |
| 23 | 3days | M17 | small | | yes | *L. nagelli* | | | | | | | | | | | | | | | | | | | |
| 24 | 3days | M17 | small | | yes | *L. nagelli* | | | | | | ESL0978 | | | | | | | | | | | | | |
| 25 | 3days | SC | big | flach | yes | *S. cervisiae* | yes | | 1.56 | | | ESL0990 | | | | | | | | | | | | | |
| 31 | 3days | FH | small | | yes | *L. nagelli* | | | 54 | | | | | | | | | | | | | | | | |
| 32 | 3days | FH | small | | yes | *L. nagelli* | | | 49 | | | | | | | | | | | | | | | | |
| 33 | 3days | FH | small | | yes | *L. nagelli* | | | 45.8 | | | | | | | | | | | | | | | | |
| 34 | 3days | FH | small | | yes | *L. nagelli* | | | 47.7 | | | | | | | | | | | | | | | | |
| 36 | 8days | M17 | kleiner | | yes | ***no hit*** | yes | | LOW | | | ESL0979/ESL0992 | | | | | | | | | | | | | |
| 37 | 8days | FH | small | transparent | yes | ***no hit*** | yes | | LOW | | | | | | | | | | | | | | | | |
| 38 | 8days | FH | small | transparent | yes | ***no hit*** | yes | | 7.58 | | | ESL0980/ESL0993 | | | | | | | | | | | | | |
| 39 | 8days | FH | big | | yes | *L. paracasei* | yes | | | | | ESL0981 | | | | | | | | | | | | | |
| 40 | 8days | FH | klein | transparent | yes | *L. nagelli* | yes | | | | | | | | | | | | | | | | | | |
| 41 | 8days | FH | big | | yes | *L. parabucherni* | yes | | | | | ESL0982 | | | | | | | | | | | | | |
| 42 | 8days | ML | small | white | yes | *S. cervisiae* | yes | | LOW | | | ESL0994 | | | | | | | | | | | | | |
| 43 | 8days | ML | small | transparent | yes | *L. paracasei* | yes | | | | | ESL0983 | | | | | | | | | | | | | |
| 44 | 8days | SC | small | | yes | *Lactobacillus gha* | yes | | LOW | | | ESL0984 | | | | | | | | | | | | | |
| 45 | 8days | SC | small | | yes | ***no hit*** | yes | | Missing | | | Missing | | | | | | | | | | | | | |
| 46 | 8days | SC | small | | yes | *L. nagelli* | | | | | | ESL0985 | | | | | | | | | | | | | |
| 48 | 8days | MRS | small | | yes | *L. nagelli* | | | | | | ESL0987 | | | | | | | | | | | | | |
| 49 | 8days | MRS | small | | yes | *L. nagelli* | | | | | | ESL0988 | | | | | | | | | | | | | |
| 51 | 8days | PY | small | | yes | Dekkera bruxelle | yes | 20 | 0.46 | | | | | | | | | | | | | | | | |
| 52 | 8days | PY | small | | yes | Dekkera bruxelle | ? | 20 | 0.346 | | | | | | | | | | | | | | | | |
| | 3days | PY | small | | No | ***no hit*** | | | | | | | | | | | | | | | | | | | |
