## Supplement method III_ DNA extraction protocol for "Strain-level and phenotypic stability contrasts with plasmid and phage variability in water kefir communities"

### DNA extraction from water kefir grains

#### Pulverization

1. Weigh 5 g of fresh or frozen kefir grains.
2. Transfer kefir grains to a 50 ml grinding jar with 25 mm ball (stainless steel) for Mixer Mill MM 400 (Retsch) and tighten the lid.
3. Freeze in liquid nitrogen, e.g. in a polystyrene foam box.
4. Fix a pair of grinding jars in the Mixer Mill 400 instrument.
5. Pulverize 1.5 min at 30 Hz.
6. Chill 15 ml and 50 ml centrifuge tubes and metallic spatula in liquid nitrogen.
7. Carefully open the grinding jar and transfer 1 g of the pulverized material to a 15 ml centrifuge tube with spatula—for DNA extraction.
8. Transfer the remaining pulverized material to a 50 ml tube and store as backup frozen at -20°C or -80°C.
9. Wash the grinding jar and ball with a sponge and a detergent; rinse with water. Soak in 0.1 M NaOH for 5 min, then, thoroughly wash with tap water; finish cleaning by spraying with ethanol and wiping out with tissue. Irradiate for 5 min in UV crosslinker (optional).

#### Crude DNA extraction

10. To the 15 ml tube containing 1 g of the pulverized kefir grains, add 2 ml of 1X lysis buffer and mix by vortexing.
11. Transfer to water-bath set to 56°C.
12. When melted, add 20 μl of proteinase K (20 mg/ml) and mix by 5 sec vortexing.
13. Incubate at 56°C for 1.5 h. Mix every 15 min.
14. Add 10 μl RNAse A (20 mg/ml) and mix by vortexing 5 sec.
15. Incubate 15 min at 37°C.
16. Divide the volume into two 1.5 ml centrifuge tubes.
17. Centrifuge at 14000 rpm for 15 min at 20°C.
18. Transfer supernatant to two 2 ml tubes (expect about 0.9 ml per tube).
19. Add 0.8 v isopropanol (e.g., 0.72 ml per 0.9 ml) and mix by vortexing 5 sec.
20. Mix and incubate for 15 min at room temperature.
21. Centrifuge at 14000 rpm for 15 min at 20°C.
22. Discard supernatant and remove liquid traces by a pipette.
    a. NOTE: safe stopping point. For storage, add 1 ml 80% ethanol and store the pelleted crude DNA at -20°C or continue with DNA purification.

#### DNA purification using Zymoclean Large Fragment DNA recovery kit

23. To each of the crude DNA pellets obtained in step 22, add 50 μl of TER buffer (10 mM TrisHCl pH 8.0, 1 mM EDTA, and 0.2 mg/ml RNAse A).
24. Incubate for 15 min at 37°C and 600 rpm in a ThermoMixer F2.0 (Eppendorf).

25. To each tube, add 3 volumes of ADB buffer (150 ul per 50 ul DNA solution) from the Zymoclean Large Fragment DNA recovery kit.
26. Incubate for 5 min at 37°C and 600 rpm in a ThermoMixer F2.0. Verify that DNA was dissolved completely; if needed help dissolving by repeated pipetting.
27. Combine two samples (total volume ca 400 ul) and apply the mixture to a column.
28. Centrifuge 2 min at 5000 rpm and 25°C and discard the supernatant.
29. Add 150 μl of ADB.
30. Centrifuge 1 min at 12000 rpm and 25°C and discard the supernatant.
31. Add 200 μl of washing buffer.
32. Centrifuge 1 min at 12000 rpm and 25°C and discard the supernatant.
33. Repeat steps 33 and 34 two more times to have three washes in total.
34. Take the column out of the recipient tube and place it into the new recipient tube.
35. Centrifuge 2 min to dry.
36. Clean with a tissue to remove any traces of liquid.
37. Place the column to a clean 1.5 ml tube and place into a ThermoMixer F2.0 set to 55°C.
38. Add 30 μl of elution buffer preheated to 55°C to the center of the column matrix.
39. Incubate 5 min at 55°C.
40. Centrifuge 1 min at 12000 rpm and 25°C.
41. Repeat steps 38-40 (in total two elutions with 30 ul).
42. Save the eluate.
43. Quantify DNA concentration with NanoDrop instrument.

#### SPRI Magnetic bead DNA purification with CleanNGS beads

44. Prepare PCR tubes in strips.
45. Per 50 μl DNA sample, add 40 μl of CleanNGS beads (Clean NA) and mix by vortexing 5 sec.
46. Incubate 5 min at RT. Every 1 min vortex to mix.
47. Place tubes on a magnet stand and do all manipulations on the magnet.
48. Incubate 5 min.
49. Pipet liquid out.
50. Add 200 μl of 80% ethanol.
51. Incubate 5 min.
52. Pipet liquid out.
53. Add 200 μl of 80% ethanol.
54. Pipet liquid out. Pay attention to removing all the ethanol.
55. Dry for 5 min in a sterile hood.
56. Add 40 μl of 5 mM Tris-HCl pH 8.0 preheated to 55°C.
57. Mix by tapping on the tube strips until homogenous.
58. Incubate 5 min.
59. Pulse-spin to bring all the liquid to the bottom.
60. Place tubes to the magnet stand.
61. Incubate 5 min (or until clear).

62. Transfer the supernatant containing DNA to a new tube.
63. Quantify DNA by NanoDrop and/or Qbit and analyze 2 μl on the agarose gel. To avoid repeated freezing-thawing, make 2 or 3 aliquots and store frozen.

#### Materials and solutions

- 1xLysis buffer: 50 mM TrisHCl pH 8.0; 25 mM EDTA pH 8.0; 1% Tween 20; 1% Triton X-100; 4 M Guanidinum HCl (304 g/l).
  *Note: high concentration of guanidinium hydrochloride was meant for direct column purification of cell lysates, e.g. in combination with Qiagen genomic tips. However, in the pilot experiments, we observed significant carryover of non-DNA material after the column purification, consequently , leading to protocol modifications described above.*
- Proteinase K 20 mg/ml
- RNAse A 20 mg/ml
- TER buffer: 10 mM Tris-HCl pH 8.0, 1 mM EDTA, 0.2 mg/ml RNAse A
- 5 mM Tris-HCl pH 8.0
- Isopropanol
- 80% ethanol
- CleanNGS DNA & RNA Clean-Up Magnetic Beads
